# Integrative construction of regulatory region networks in 127 human reference epigenomes by matrix factorization

**DOI:** 10.1101/217588

**Authors:** Dianbo Liu, Jose Davila-Velderrain, Zhizhuo Zhang, Manolis Kellis

## Abstract

Despite large experimental and computational efforts aiming to dissect the mechanisms underlying disease risk, mapping cis-regulatory elements to target genes remains a challenge. Here, we introduce a matrix factorization framework to integrate physical and functional interaction data of genomic segments. The framework was used to predict a regulatory network of chromatin interaction edges linking more than 20,000 promoters and 1.8 million enhancers across 127 human reference epigenomes, including edges that are present in any of the input datasets. Our network integrates functional evidence of correlated activity patterns from epigenomic data and physical evidence of chromatin interactions. An important contribution of this work is the representation of heterogeneous data with different qualities as networks. We show that the unbiased integration of independent data sources suggestive of regulatory interactions produces meaningful associations supported by existing functional and physical evidence, correlating with expected independent biological features.

## Introduction

The disruption of cis-regulatory elements is considered the key mechanism through which disease risk is conferred by noncoding mutations (1–3). However, in order to support this hypothesis and apply it in the development of rational therapeutic strategies, several difficulties have to be surpassed. First, identification of cis-regulatory elements proved a difficult task given the dimension of the noncoding genome (4). This has been overcome using the association of chromatin marks with genome activity in coding and noncoding regions as a widely accepted approximation to map the tissue-specific activity and dynamics of distal and proximal cis-regulatory elements (5–7). The highly correlated structure displayed by combinatorial patterns of marks across the genome enables computational identification of a reduced number of robust chromatin states (8,9) for display in a single annotation track. Thanks to these experimental and computational advances, reference epigenomes were recently profiled and annotated for a large number of human tissues (10), including the tissue-specific annotation of active cis-regulatory elements (e.g., enhancers).

Having defined systematic strategies for genome-wide mapping of cis-regulatory elements, efforts have more recently shifted towards tackling the more challenging problem of determining what genes are likely to be targeted by given cis-regulatory elements, mostly enhancers. Numerous solutions have been proposed on both the computational and experimental fronts.

On the computational side, several efforts have exploited the correlated structure of epigenomic features to infer associations between enhancers and target promoters. Enhancer-promoter associations have been mapped by quantifying patterns of coactivity of annotated enhancer elements and promoters across and within tissues (8,10). Supervised machine learning approaches with the goal of learning epigenomic patterns discriminative of functional interactions have also been proposed (11–13). On the experimental side, techniques to measure chromatin conformation enable the mapping of high confidence interactions at different levels of resolution and across several human cell-types and tissues (14,15), These methods can be targeted to elucidate regulatory interactions by enrichment of potential enhancer-promoter contacts in assays like the Chromatin Interaction Analysis by Paired-End Tag Sequencing (ChIA-PET) or promoter capture Hi-C, a promoter centered chromosome conformation capture technique (14,16). However, both approaches suffer from limitations. First, there is no gold standard interaction set. Second, it is currently not feasible to profile chromatin interactions in a large number of cells and tissues to provide a tissue-specific reference for an organism. In addition, the level of resolution of Hi-C experiments makes it far from trivial to precisely localize the particular enhancer and promoter pairs that might be involved in functional transcriptional regulatory interactions. Given these and associated limitations, and the availability of recently published human reference epigenomes (127 cell/tissue types) (10) and the largest sets of mapped chromatin interactions across human tissues (17 primary blood cell types and 21 cell/tissue types) (14,15), we reasoned that a hybrid and integrative computational approach is timely.

This article presents SWIPE-NMF, a computational method that implements Sliding WIndow PEnalized Nonnegative Three-factor Matrix Factorization on heterogeneous association data represented as networks. This approach was used to integrate the functional and physical evidence of regulatory interactions provided by computational coactivity inference and experimental data, respectively. This method was applied to annotate a weighted set of potential interactions for each of the 127 cell and tissue types within human reference epigenomes (10). Furthermore, SWIPE-NMF was implemented as a flexible tool that can be applied to integrate any set of enhancer annotations with prior evidence sources for regulatory interactions to infer tissue-specific weighted interactions.

## Results

### A matrix dimensionality reduction framework integrating evidence of enhancer-promoter interactions

In order to computationally integrate the large set of qualitatively different data suggestive of potential enhancer-promoter interactions in a principled way, we first curated a database including five experimental data sources (Figure 1). We considered previously published (1) enhancer-promoter coactivity associations(8,10), (2) physical chromatin interaction calls from Hi-C data in 21 tissues (15,17), (3) cis-regulatory-gene associations defined by eQTL in 53 human tissues (18), (4) cis-regulatory-promoter associations defined by activity correlation between DNase-I hypersensitivity sites (DHS) and promoters (19), and (5) topologically associated domain (TAD) annotations defined from Hi-C data (20). A description of each data type and the nature of evidence provided is included in the *Supplementary material* (S Table 1). As an example, Figure 1A shows the density of data in a randomly selected region in K562 cells.

**Figure 1.**
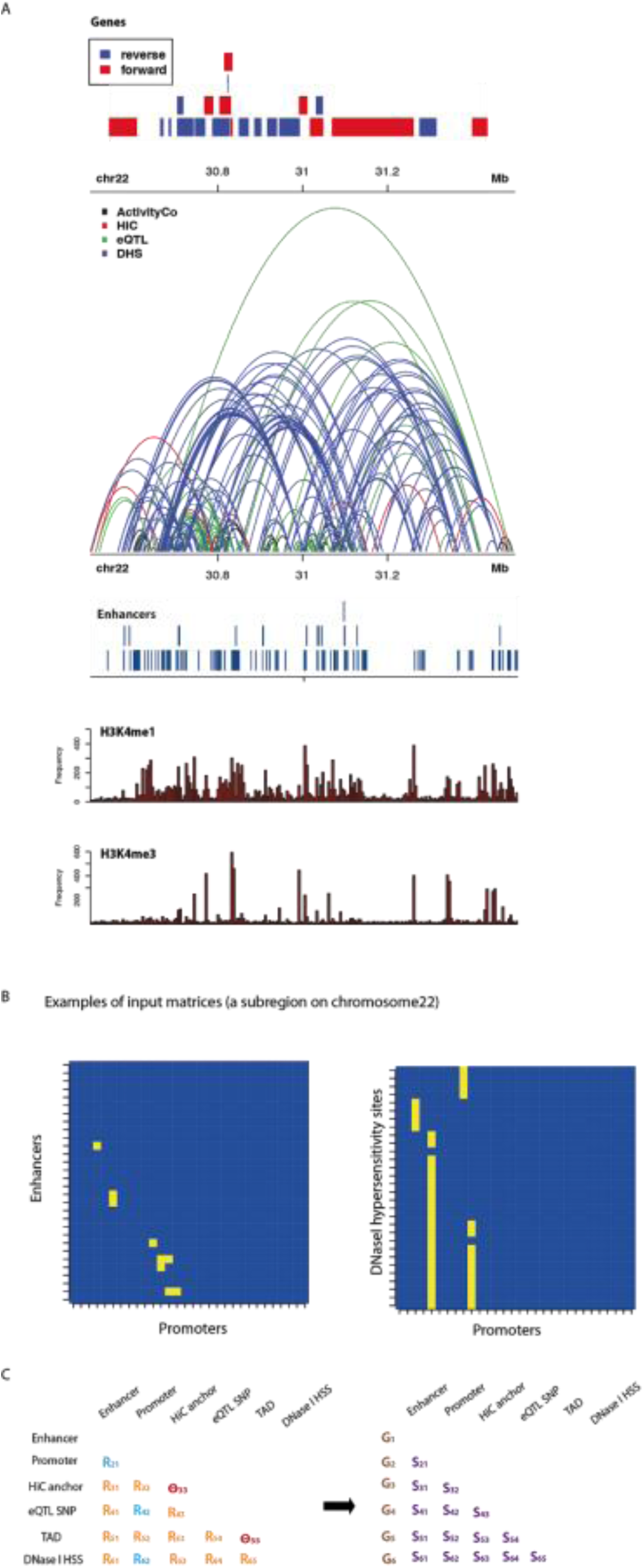
Schematic representation of the SWIPE-NMF framework. Heterogeneous association data coded as binary networks were integrated, and scored sets of tissue-specific enhancer-promoter, enhancer-enhancer, and promoter-promoter interactions inferred for 127 human reference epigenomes in an unsupervised manner. **A)** Different genetic interactions in a randomly selected region on chromosome 22. Enhancer-promoter activity correlations (EP), are shown in blue. Hi-C links are in red. Links between SNPs and promoters detected by eQTL are in green. Correlation between DNaseI hypersensitivity sites and promoters across multiple cell types are in sky blue. Topologically associated domains are not shown to avoid confusion with links among genomic elements. Locations of genes, reference enhancers and histone marks were also included. **B)** All data types were organised into a matrix/networks. Each row or column represents a type of genomic segments such as enhancers, promoters or Hi-C anchors. **C)** SWIPE-NMF was used to integrate all five data types to produce cell/tissue types. Each matrix *R*_*ij*_ was decomposed into three matrices *G*_*i*_, *S*_*ij*_ and 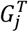 such that 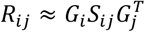. *R*_*ij*_ is the relation between the data tyes *i* and *j*. *R*_12_ is enhancer promoter interaction. *G*_*i*_ is an *n* × *m* matrix where *n* is the number of elements in that data type (e.g., number of enhancers) and *m* is the number of ranks. *S*_*ij*_ is a matrix representing the relation between columns in *G*_*i*_ and *G*_*j*_ Joint factorization of matrices allows integration of information from all data types while minimizing information loss. This factorization was conducted on 5 Mb overlapped windows on each chromosome in each cell and tissue type.

For consistency, we defined a reference set of potential enhancer elements to which all data were mapped. The reference selected was the non-genic enhancer ChromHMM chromatin state (7-Enh) annotated for all 127 reference human epigenomes in the roadmap epigenomics project (10). The heterogeneous nature of the data, i.e., association data at different length scales, in addition to annotations of discrete genomic regions (e.g., TAD domains), made the integration task challenging. We approached the problem by first devising an individual network representation for each data source representable in matrix form and compatible for mapping across sources (Figure 1B, C). We then applied an extended NMF algorithm to fuse the independent network data.

Specifically, we considered six types of genomic segments: enhancer, promoter, Hi-C anchor, cis-eQTL (i.e., the SNP position having the association), DHS, and TAD. Each network is composed of the total genomic segments in the data. We can define two qualitatively different types of associations (Figure 1C): interaction and incidence associations. Interaction matrices (blue) code associations between genomic segments of different types that are supported by either physical or functional experimental data. Incidence matrices represent the incidence of one element within the other (orange) i.e, one genomic segment overlapping with the other. Finally, we define two diagonal incidence matrices ϴ (red), which operationalize the prior knowledge that regulatory interactions are expected to be supported by Hi-C physical interactions and preferentially occur within TAD domains. We defined a consistent set of matrices for each cell/tissue type (for details, see Materials and Methods). Thus, we solved the problem of heterogeneous data representation by operationally defining networks of experimentally supported association as binary matrices *R*_*ij*_ that code associations between genomic segments of type *i* and *j*. Importantly, integrating the data into this format enables the application of well-established matrix factorization algorithms. The matrices *R* and ϴ are the inputs of the matrix factorization algorithm.

### Three-factor penalized matrix factorization (PMF)

Our method extends the traditional three-factor penalized matrix factorization (PMF) approach, which has been recently used for gene functions and pharmacologic actions predictions with an additional constraint imposing genomic locality of regulatory interactions (21–23). This method is designed to fuse the heterogeneous network datasets and infer a scored set of enhancer-promoter, enhancer-enhancer, and promoter-promoter interactions (Figure 1C).

Our method seeks to decompose the observed interaction matrix into a lower-dimensional representation that reveals biologically-meaningful components. All the association matrices *R*_*ij*_ are simultaneously factorized: each individual matrix is decomposed into *G*_*i*_, *G*_*j*_ and *S*_*ij*_ so that 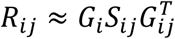. In other words, an entry *R*_*ij*_(*p*,*q*) is approximated by the inner product of the *p*-th row of matrix *G*_*i*_ and a linear combination of the columns of *S*_*ij*_, weighted by the *q*-th column of matrix. *G*_*i*_. The objective function to minimize is:

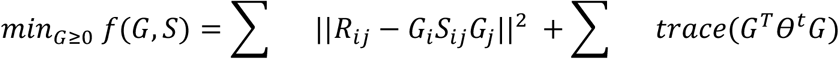

From a biological perspective, a matrix R_ij_ defines the association between two different genomic segment types *i* and *j*, such as enhancers and promoters. A matrix *G* is specific to a type of genomic segments and records associations among genomic segments of that type.

- Each row of *G*_*i*_ is a genomic segment of type i (e.g., an enhancer).
- The columns of *G*_*i*_ can be understood as clusters dividing genomic segments of type *i* based on shared patterns of regulatory or physical interactions.
- The matrix *G*_*i*_ specifies the probability of each genomic segment of type *i* belonging to each cluster.
- The matrix *S*_*ij*_ can be interpreted as defining association among clusters of genomic segment type *i* and type *j*.

Using a sliding window of size 5Mb, the matrix factorization algorithm focuses on uncovering local patterns within the windowed genomic region (see Materials and Methods for details). Association among genomic segments of the same type, such as enhancer-enhancer interactions, can be estimated from *G*_*i*_ matrices by *G*_*i*_*G*_*j*_^*T*^. In this way, *R*_*ij*_ is dissected into, and can be reconstructed from, three matrices,*G*_*i*_, *G*_*j*_ and *S*_*ij*_ in a systematic, tractable and interpretable way.

The algorithm iteratively updates G and S by fixing one of them in an alternate way. We applied this method to all the integrative tissue-specific sets of matrices, including Hi-C, enhancer-promoter activity correlation, DHS-promoter correlation, eQTL, and TAD domains; obtaining a tissue-specific weighted set of interaction matrices for enhancer-promoter (reconstructed *R*_12_), enhancer-enhancer(*G*_1_*G*_1_^*T*^), and promoter-promoter(*G*_2_*G*_2_^*T*^) interactions.

### Evaluation of the data integration strategy

There is currently no large gold-standard compendium of known regulatory region interactions, and lines of evidence for physical, functional, and genetic interactions each capture different aspects of the underlying regulatory network. However, these complementary biological datasets enabled us to validate our predictions using a diversity of methods and evidence. (1) For enhancer-promoter co-activity associations, we used five-fold cross validation; (2) for each independent empirical data set suggestive of regulatory associations, we excluded one whole dataset from inputs; and (3) with orthogonal experimental data, we used separate ChiA-PET data. Finally, (4) we examined biological correlates and cell/tissue-type specificity of the scored sets of interactions inferred by SWIPE-NMF.

In five-fold cross validation, SWIPE-NMF was used to reconstruct the functional coactivity data of enhancer-promoter associations (8), with 20% of the associations excluded from inputs. SWIPE-NMF showed good performance on this task (AUC = 0.82) (Fig. 2A). Next, in four evaluation experiments, each evidence source (HiC, eQTL, TAD and DHS) was left out and used to test the model’s consistency with the interactions inferred by integrating the rest of the datasets. When each eQTL and DHS was individually excluded from inputs and used as ground truth, the inference also performed well (AUROC > 0.7, Fig 2B), indicating that the interactions predicted by integration are supported by eQTL and DHS correlation (19). In addition, when TAD incidence annotation is excluded from inputs, the corresponding inferred interactions occurring within TAD domains have much higher scores compared with those involving cross-domain interactions (p-values <10^−10^) (Figure 2C). Finally, most of the interactions with high confidence scores are within TAD domains (15)(Figure 2D). When testing using orthogonal ChiA-PET data (from the K562 cell line) (24), the performance of SWIPE-NMF (AUROC > 0.70) is better compared with either enhancer promoter activity correlation (AUROC≈0.6), averaging links across data types (AUROC≈0.6), and a simple nearest promoter assignment (the brown point in Figure 2E). The AUROC is different from previous publications because enhancers are defined in different way, we only focus on enhancers in non-genic regions in this project and input data were handled in a more conservative way (see Materials and Methods for details). Interestingly, we found that a considerable portion (50% to 80% depending on cell and tissue type) of enhancer-promoter interactions inferred by SWIPE-NMF with low scores do not occur in any of the input datasets (Figure 2F). When comparing with ChiA-PET links, we found that these interactions uniquely predicted by SWIPE-NMF show better performance than random expectation (AUROC≈0.6). This suggests factorization is able to transfer information by learning association patterns in observed enhancer-promoter interactions. The results are consistent with orthogonal data of chromatin interactions mediated by RNA polymerase (24). Presumably the degree of overlap will increase when matching tissue-specific ChiA-PET data are considered, once available. Overall, the evaluation experiments demonstrate that, through integration by SWIPE-NMF, different sources of evidence provide complementary information with predictive power.

**Figure 2.**
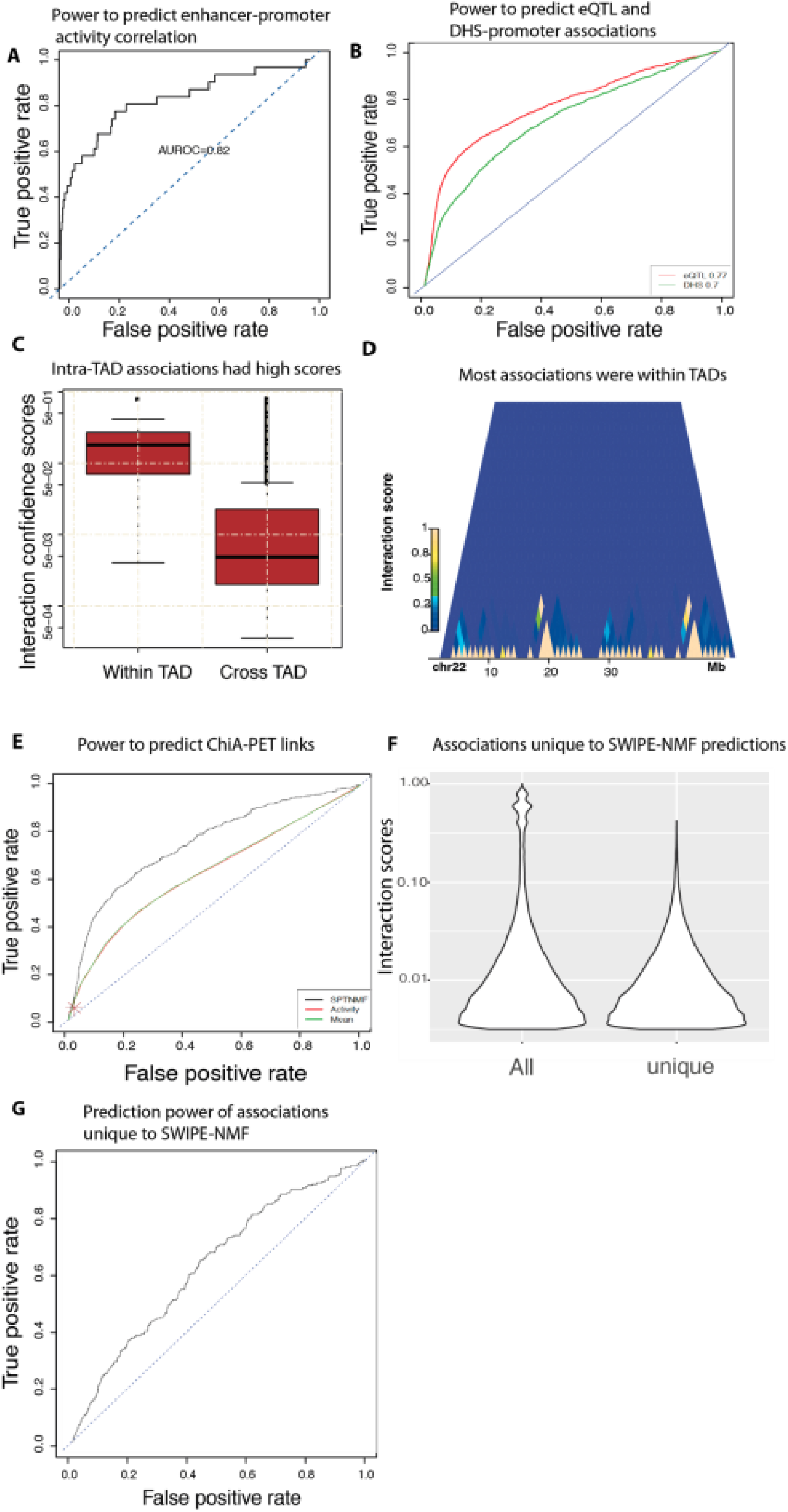
Performance of SWIPE-NMF in enhancer-promoter interaction inference. A) receiver operating characteristic (ROC) curve was used to demonstrate the power of SWIPE-NMF method to reconstruct enhancer-promoter network inferred by activity correlation alone (five-fold cross validation). 20% of the enhancer-promoter activity correlation links were left out in each fold. An area under ROC curve (AUROC) > 0.8 was reported (AUORC≈0.5 for random predictions). B) Performance of SWIPE-NMF by leaving each of the entire datasets out in inputs. Using eQTL-promoter links (red), and DNaseI hypersensitivity site to promoter correlation links (green) as ground truth, AUROC were both > 0.70. C) Confidence scores of interactions within topologically associating domains (TAD) are significantly higher than inter-TAD interactions with a P value *<10*^*−5*^. D) Most of the high score interactions are within TAD domains. Each block on x axis is a TAD. Chromosome 22 of the K562 cell line is shown. Yellow color indicates high interaction scores and blue color indicates lower scores. E) Performance of SWIPE-NMF using ChiA-PET as gold standard (24). SWIPE-NMF (black, AUROC>0.7) performs better than activity based correlation (red, AUROC≈ *0.6*), average matrix (green, AUROC≈ *0.6*), and nearest promoter (brown). F) 50–80% of the enhancer-promoter links were unique to output of SWIPE-NMF, i.e., not seen in any of the five input data types. The confidence scores of links unique to SWIPE-NMF were generally in middle to lower range. G) Enhancer-promoter links unique to SWIPE-NMF output had an AUROC≈ *0.6* when ChiA-PET was used as ground truth.

### Predicted enhancer-promoter interactions are biologically meaningful

Several studies have shown a strong correlation between chromatin interactions and gene co-expression, due to the spatial colocalization of transcribed genes and their regulatory elements (25,26). We tested whether the inferred associations present a similar behavior. Using a large set of tissue-specific gene coexpression networks (27), we found that coexpressed gene pairs tend to share common interacting enhancers (P-val < 1e-30, hypergeometric test), agreeing with the expected behavior (Figure 3A). We also found enrichment of transcription factor binding sites (TFBSs) within enhancers and promoters, and moreover, we show that inferred interacting enhancer-promoter pairs sharing TF binding motifs are more likely to interact (Figure 3B) than those without co-occurring motifs. These results are consistent with previous reports suggesting that transcription factors might facilitate enhancer-promoter interactions (27,28). Previous studies have also shown that CTCF, an insulator binding protein that is thought to be involved in the regulation of chromatin structure and DNA looping (29), is enriched near interacting promoters and enhancers (11,13). We found enhancers interacting with promoters and promoters interacting with enhancers are both highly enriched in CTCF Chip-seq peaks with significant p-values (Figure 3 C and D), within the inferred associations.

**Figure 3.**
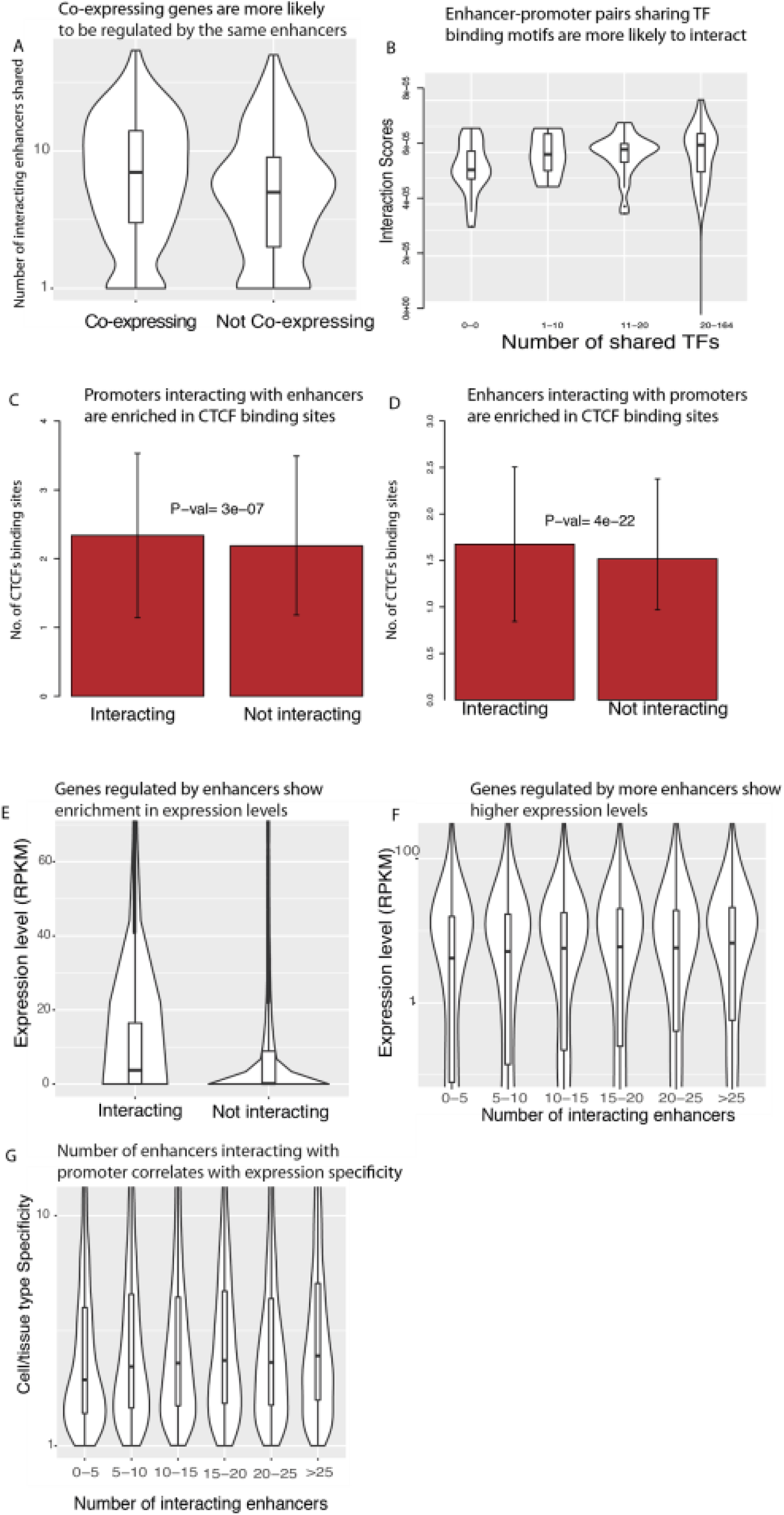
Biological correlates of inferred enhancer-promoter networks produced. No enhancer-enhancer and promoter-promoter links are considered in this figure. A) Co-expressing gene pairs are more likely to share interacting enhancers compared with gene pairs not showing co-expression (P-val < *10*^*−15*^) (27). Data for K562 cell was shown. B) Enhancer-promoter pairs sharing more TF binding motifs are more likely to interact(p value<*10*_*−15*_). C) Promoters interacting with enhancers show enrichment in CTCF binding sites detected by Chip-Seq which agrees with previous findings (13) D) Enhancers interacting with promoters also show enrichment in CTCF binding sites detected by Chip-Seq. E) Genes regulated by enhancers show enrichment in expression levels compared with genes not interacting with enhancers. F) Numbers of interacting enhancers of genes have a positive correlation with expression levels. G) Number of enhancers interacting with each promoter also positively correlates with cell/tissue type expression specificity measured by entropy rank (11).

### Tissue-specific promoters show more enhancer interactions

Enhancers are known to regulate tissue-specificity predominantly by modulating the expression of different target genes across tissues (30). We tested whether genes being targeted by enhancers show distinctive properties of gene expression relative to other genes using the inferred interactions. We found that genes interacting with enhancers show higher expression (RPKM) than those without enhancer regulation (Figure 4.3E). In addition, the level of gene expression in each cell type shows a positive correlation with the number of incoming enhancer interactions to the promoters, consistent with an additive effect of the regulatory input (Figure 3F). Furthermore, given the role of enhancers in establishing tissue-specificity, we hypothesized that genes with tissue-specific functionality are more prone to targeting by enhancers. To test this hypothesis, we used an entropy based measure of gene expression specificity for each gene across the reference human transcriptome of the Roadmap epigenomics project (10,11), and found that gene expression specificity does correlate with the number of incoming enhancer interactions of a promoter (Fig. 3G). This result is consistent with the expectation that enhancer-promoter interactions contribute to the cell/tissue type specific expression of genes and agrees with findings in previous publications (11).

**Figure 4.**
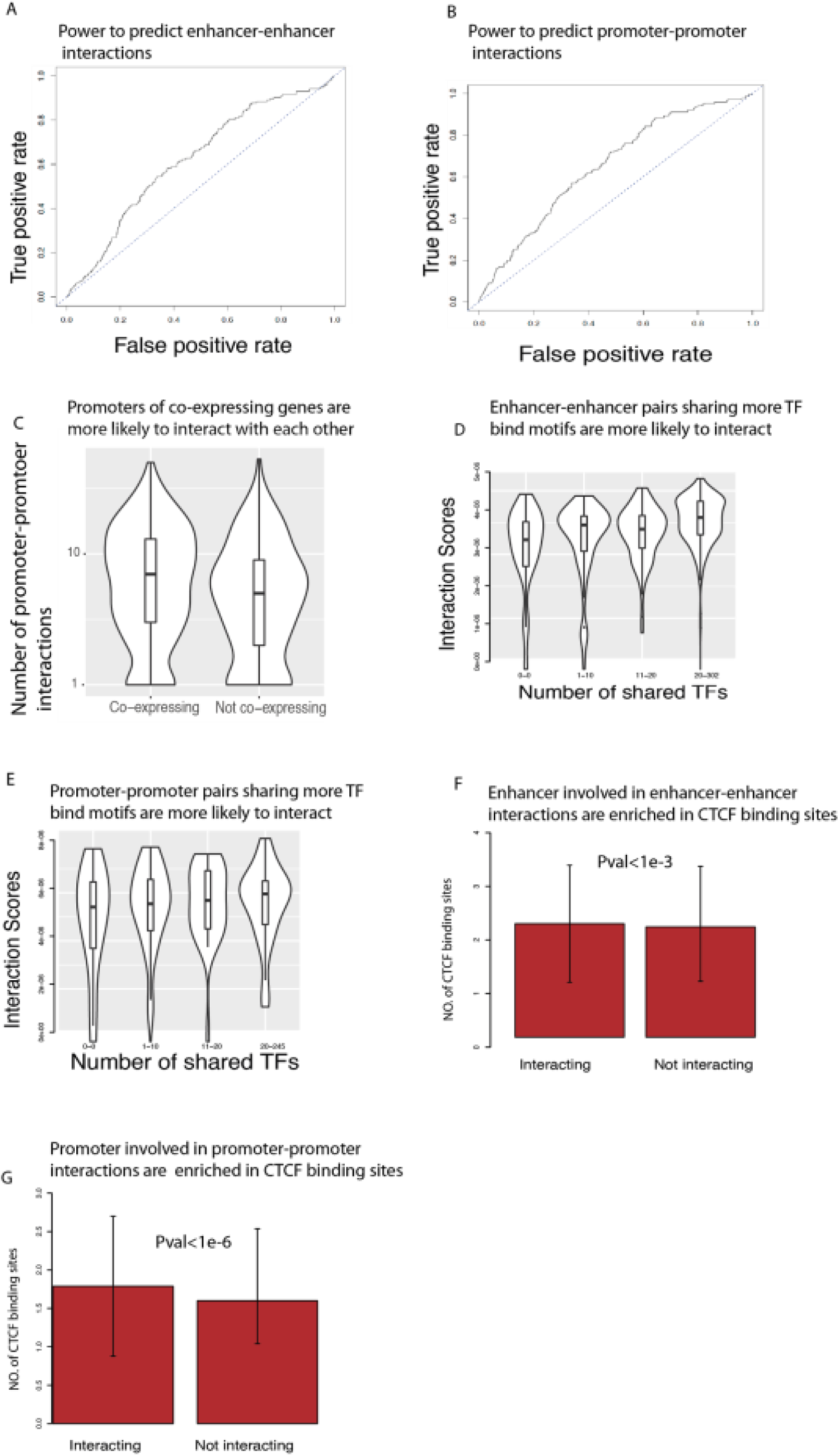
Biological properties of enhancer-enhancer and promoter-promoter interactions are similar to enhancer-promoter links. A,B) ROC curve of enhancer-enhancer and promoter-promoter interactions predicted by SWIPE-NMFusing ChiA-PET as gold standard both >0.6. Results of K562 are shown. C) Promoters of co-expressing genes are more likely to interact with each other. D,E) Enhancer-enhancer pairs and promoter-promoter pairs sharing more TF bind motifs are more likely to interact. F,G) Enhancers involved in enhancer-enhancer and promoters involved in promoter-promoter interactions are enriched in CTCF binding sites detected by Chip-Seq.

In order to further test whether genes targeted by more enhancers tend to be associated with tissue-specific (related) functions, we performed gene ontology (GO) term enrichment tests using the genes with the top 5% of incoming enhancer interactions as gene query set. In Table 5.1, we show examples of the enriched terms found for randomly chosen cell and tissue types. By looking at the top 3 GO terms in biological processes for each cell and tissue types, we found that highly targeted genes were generally enriched in functions related to the underlying biology of the tissue. This further supports our hypothesis that the inferred enhancer-promoter interactions contribute to the regulation of tissue-specificity.

**Table 5.1.**
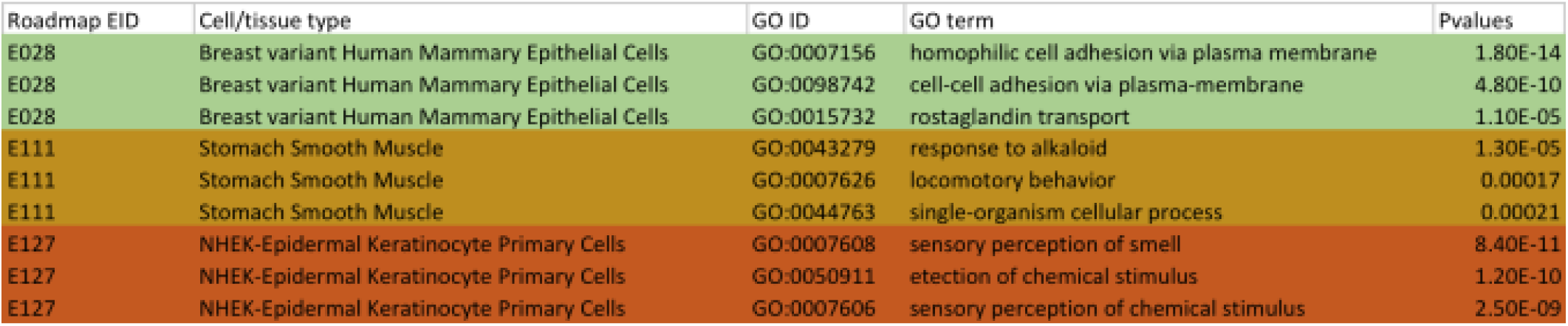
Gene ontology term enrichment

### SWIPE-NMF enhancer-promoter, enhancer-enhancer, and promoter-promoter interactions

In addition to enhancer-promoter interactions, chromatin interactions involving only promoters or only enhancers have been shown to spatially organize the transcriptional machinery (24,31). Although the mapping and characterization of enhancer-promoter interactions has received much more attention, chromatin interactions occurring at similar resolution but involving only promoters or enhancers might be relevant under normal and abnormally disrupted conditions. One advantage of using SWIPE-NMF for data integration is that all three types of chromatin interactions are simultaneously learned during the matrix factorization process (see details in Materials and Methods). When using ChiA-PET as gold standard, both enhancer-enhancer and promoter-promoter networks show considerable AUROC scores (>0.6) (Fig. 4A, B). Similar to enhancer-promoter interactions, promoters interacting with each other tend to preferentially show co-expression (Fig. 4C), consistent with the existence of chromatin mediated transcription factories within the cell(32). Interacting enhancer-enhancer pairs and promoter-promoter pairs sharing more TF motifs have higher chances of interaction (Figure 4D and 4E), and interacting enhancers and promoters are enriched in CTCF ChIP-Seq peaks (Figure 4F and 4G). Overall, the observed consistency of biological correlates across the different types of inferred interactions indicates, by integrating physical and coactivity evidence of association, SWIPE-NMF is able to infer general chromatin interactions, with enhancer-promoter associations as an important subset.

## Discussion: construct enhancer promoter networks using an intermediate integration strategy

After mapping cis-regulatory elements to their target genes, testable mechanistic hypotheses can be proposed for detrimental effects conferred by non-coding pathogenic mutations. With the goal of accelerating such mapping genome-wide, and to provide a starting reference set of potential chromatin mediated regulatory interactions across reference human tissues, here we introduced and applied SWIPE-NMF.

Several features distinguish the proposed computational framework from other tools concerned with particular instances of the same problem. When dealing with data integration, most existing methods either transform all data sources into a single feature-based table and apply to it well-established feature-based machine learning algorithms (*early integration)*, or build an independent model for each data source (*late integration)*. SWIPE-NMF, instead, is based on a more recent, *intermediate integration* strategy that explicitly addresses the multiplicity of data types by fusing them through inference of a single joint model (33,34). Importantly, such an intermediate level of integration retains the structure of the data sources, incorporating them within the structure of the learned model. SWIPE-NMF was specifically designed to exploit the information provided by both computational coactivity-based inferences and experimentally grounded physical evidence of chromatin interactions, overcoming their individual limitations. An important contribution of this method is the representation of heterogeneous data with multiple resolutions as networks, enabling their integration without resolution conversion. SWIPE-NMF implements an unbiased, unsupervised approach that directly factorizes all the integrated data matrices using non-negativity constraints (35). The simultaneous factorization of matrices allows sharing of information by revealing the latent structure of all input network data. Finally, SWIPE-NMF is applied using overlapped sliding windows along chromosomes, facilitating the capture of both local and global patterns from the data, while at the same time improving efficiency. Although, in this project, five experimental datasets were selected to provide reliable resources of interactions among promoter and enhancers, the proposed framework is flexible and can easily take into account other datasets of interest. Moreover, the method can be applied to other purposes such as improving predictions of eQTLs and chromatin physical interactions.

This new matrix factorization based approach integrates independent data sources suggestive of regulatory interactions. Application to a large set of reference human tissues produces meaningful associations supported by existing functional and physical evidence, and which correlate with expected, independent biological features. The integrative emphasis underlying the design of our approach limits its predictive reach, as the quality and quantity of inferred interactions depends on the status of the available data. However, we consider this as a strength of our approach on inferred sets of tissue-specific interactions. Data are being produced and curated at an accelerated pace. By integrating new data, SWIPE-NMF will enable inference of novel associations and improvement of the current analysis. Unbiased and integrative computational tools are required to fully exploit the power of the multiple flavors of next-generation sequencing data and epigenomic information.

## Materials and Methods

### Datasets and data processing

High resolution HiC interactions were obtained for a total of 21 human cell lines and primary tissues (15,17); in both cases, the significant interactions reported by the authors were used. Cell/tissue types were matched to the corresponding reference epigenome identifiers (EID) from the Roadmap epigenomics project, or matched to the closest EID according to information from the authors. Only interactions with q-value <1e-3 were considered. Tissue-specific coactivity based enhancer-promoter associations inferred as described in Ernst *et al.* 2011 were obtained from the Roadmap Epigenomics Consortium 2015(10). eQTL data (V6p) was obtained from the GTEX consortium, considering only associations with a p-value < 1e-5. TAD annotations were obtained from Dixon *et al.* 2012(20). DHS data was obtained from Thurman *et al.* 2012(19), considering only associations with a score > 0.9. Transcription factor binding motifs were obtained from Marbach *et al.* 2016(36). ChIA-PET data were obtained from Li *et al*. 2012(24). The tissue-specific enhancer annotations used as reference were extracted from the Roadmap epigenomics project, using the non-genic chromatin state (7_Enh) annotated with the core ChromHMM 15-state model. CTCF binding peaks were downloaded from ENCODE website; Broad and Narrow peaks were combined (4).

### Three factor penalised matrix factorization

#### Tissue-specific input preparation

Tissue-specific enhancer elements and enhancer-promoter coactivity associations were used for all tissues, and matched Hi-C and eQTL data when available. When not available, the union of the latter over the total available cell and tissue types was used as global reference of potential association. Likewise, global, tissue-agnostic DNAse-I activity associations and TAD domain annotations were considered for all tissues.

#### Sliding-window factorization

The algorithm used to conduct the matrix factorization was adapted from the original method proposed by Žitnik & Zupan 2015(21) and modified with sliding window settings and slightly different matrix operation algorithms. Factorization was conducted on sliding windows of 5M bp with 50% overlap along each chromosomes to focus on local patterns and at the same time make the optimization task computationally feasible. Different values for the hyperparameter *k* were used. The rank of each matrix is integer *N*k*, where *N* is the number of columns in the data type. 10 different values of *k* (0.05,0.1,0.15,0.2,0.25,0.3,0.35,0.4,0.45,0.5) were tried on each window. As different initialization of G matrices give different factorization and there is no guarantee of global minimum, we used an ensemble learning strategy of running 20 rounds of the algorithm with slightly different initializations and averaged the outputs (21). The rank for each window was determined by selecting *k* where a maximum kink was attained in total reconstruction error curve (21,37). The algorithm was stopped if the difference between two iterations was smaller than 0.01 or the maximum number of interaction (200) was reached.

#### Enhancer-promoter interaction set

In order to provide a set of high-confidence scored interactions, in addition to the direct output from the inference, we determined a cut-off value for the enhancer-promoter association score produced by SWIPE-NMF, filtering out interactions with lower scores. The cutoff was set so that the average number of promoters per enhancer was consistent with previous estimates (~3) (11). Given that a similar criterion for enhancer-enhancer and promoter-promoter interactions is not available, the filtering step was not performed for these.

#### Enhancer-Enhancer and promoter-promoter interactions

The factor *G*_2_ (Figure. 5.1C) produced by three-factor penalised factorization provides information about the learned structures of promoter networks. A weighted enhancer-enhancer interaction matrix was calculated as 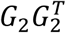 for each sliding window. Similarly, an enhancer-enhancer interaction matrix was calculated as *G*_1_*G*_1_^*T*^.

### Performance evaluation

For five-fold cross validation experiments, 20% of the associations were randomly chosen and excluded from inputs. In addition, an equal number of non-interacting pairs were randomly selected to balance the data. When a whole dataset was left out to evaluate the performance of SWIPE-NMF, that dataset was used a ground truth for testing. Only interactions occurring at a distance larger than 5kb were considered for the analyses of biological correlates.

### Availability

All the computational scripts are available at https://github.com/kaiyuanmifen/SWIPE-NMF

All the regulatory region networks are available from Broad Institute server and will deposited in online repository.

## References

1. Ward, L.D. and Kellis, M. (2012) Interpreting noncoding genetic variation in complex traits and human disease. Nat. Biotechnol., 30, 1095–1106.

2. Deplancke, B., Alpern, D. and Gardeux, V. (2016) The Genetics of Transcription Factor DNA Binding Variation. Cell, 166, 538–554.

3. Gandal, M.J., Leppa, V., Won, H., Parikshak, N.N. and Geschwind, D.H. (2016) The road to precision psychiatry: translating genetics into disease mechanisms. Nat. Neurosci., 19, 1397–1407.

4. Consortium, E.P. (2012) An integrated encyclopedia of DNA elements in the human genome. Nature, 489, 57–74.

5. Shen, Y., Yue, F., McCleary, D.F., Ye, Z., Edsall, L., Kuan, S., Wagner, U., Dixon, J., Lee, L., Lobanenkov, V.V. et al. (2012) A map of the cis-regulatory sequences in the mouse genome. Nature, 488, 116–120.

6. Vermunt, M.W., Reinink, P., Korving, J., de Bruijn, E., Creyghton, P.M., Basak, O., Geeven, G., Toonen, P.W., Lansu, N., Meunier, C. et al. (2014) Large-Scale Identification of Coregulated Enhancer Networks in the Adult Human Brain. Cell Rep., 9, 767–779.

7. Villar, D., Berthelot, C., Aldridge, S., Rayner, T.F., Lukk, M., Pignatelli, M., Park, T.J., Deaville, R., Erichsen, J.T., Jasinska, A.J. et al. (2015) Enhancer evolution across 20 mammalian species. Cell, 160, 554–566.

8. Ernst, J., Kheradpour, P., Mikkelsen, T.S., Shoresh, N., Ward, L.D., Epstein, C.B., Zhang, X., Wang, L., Issner, R., Coyne, M. et al. (2011) Mapping and analysis of chromatin state dynamics in nine human cell types. Nature, 473, 43–49.

9. Ernst, J. and Kellis, M. (2012) ChromHMM: automating chromatin-state discovery and characterization. Nat. Methods, 9, 215–216.

10. Roadmap Epigenomics, C., Kundaje, A., Meuleman, W., Ernst, J., Bilenky, M., Yen, A., Heravi-Moussavi, A., Kheradpour, P., Zhang, Z., Wang, J. et al. (2015) Integrative analysis of 111 reference human epigenomes. Nature, 518, 317–330.

11. He, B., Chen, C., Teng, L. and Tan, K. (2014) Global view of enhancer-promoter interactome in human cells. Proc. Natl. Acad. Sci. U. S. A., 111, E2191–2199.

12. Roy, S., Siahpirani, A.F., Chasman, D., Knaack, S., Ay, F., Stewart, R., Wilson, M. and Sridharan, R. (2016) A predictive modeling approach for cell line-specific long-range regulatory interactions. Nucleic Acids Res., 44, 1977–1978.

13. Whalen, S., Truty, R.M. and Pollard, K.S. (2016) Enhancer–promoter interactions are encoded by complex genomic signatures on looping chromatin. Nat. Genet., 48, 488–496.

14. Javierre, B.M., Burren, O.S., Wilder, S.P., Kreuzhuber, R., Hill, S.M., Sewitz, S., Cairns, J., Wingett, S.W., Várnai, C., Thiecke, M.J. et al. (2016) Lineage-Specific Genome Architecture Links Enhancers and Non-coding Disease Variants to Target Gene Promoters. Cell, 167, 1369–1384.e1319.

15. Schmitt, A.D., Hu, M., Jung, I., Xu, Z., Qiu, Y., Tan, C.L., Li, Y., Lin, S., Lin, Y., Barr, C.L. et al. (2016) A Compendium of Chromatin Contact Maps Reveals Spatially Active Regions in the Human Genome. Cell Rep., 17, 2042–2059.

16. Fraser, J., Williamson, I., Bickmore, W.A. and Dostie, J. (2015) An Overview of Genome Organization and How We Got There: from FISH to Hi-C. Microbiol. Mol. Biol. Rev., 79, 347–372.

17. Rao, S.S.P., Huntley, M.H., Durand, N.C., Stamenova, E.K., Bochkov, I.D., Robinson, J.T., Sanborn, A.L., Machol, I., Omer, A.D., Lander, E.S. et al. (2014) A 3D map of the human genome at kilobase resolution reveals principles of chromatin looping. Cell, 159, 1665–1680.

18. Consortium, G.T. (2015) Human genomics. The Genotype-Tissue Expression (GTEx) pilot analysis: multitissue gene regulation in humans. Science, 348, 648–660.

19. Thurman, R.E., Rynes, E., Humbert, R., Vierstra, J., Maurano, M.T., Haugen, E., Sheffield, N.C., Stergachis, A.B., Wang, H., Vernot, B. et al. (2012) The accessible chromatin landscape of the human genome. Nature, 489, 75–82.

20. Dixon, J.R., Selvaraj, S., Yue, F., Kim, A., Li, Y., Shen, Y., Hu, M., Liu, J.S. and Ren, B. (2012) Topological domains in mammalian genomes identified by analysis of chromatin interactions. Nature, 485, 376–380.

21. Žitnik, M. and Zupan, B. (2015) Data Fusion by Matrix Factorization. IEEE Trans. Pattern Anal. Mach. Intell., 37, 41–53.

22. Gligorijević, V., Janjić, V. and Pržulj, N. (2014) Integration of molecular network data reconstructs Gene Ontology. Bioinformatics, 30, i594–600.

23. Hwang, T., Atluri, G., Xie, M., Dey, S., Hong, C., Kumar, V. and Kuang, R. (2012) Co-clustering phenome–genome for phenotype classification and disease gene discovery. Nucleic Acids Res., 40, e146–e146.

24. Li, G., Ruan, X., Auerbach, R.K., Sandhu, K.S., Zheng, M., Wang, P., Poh, H.M., Goh, Y., Lim, J., Zhang, J. et al. (2012) Extensive promoter-centered chromatin interactions provide a topological basis for transcription regulation. Cell, 148, 84–98.

25. Schoenfelder, S., Sexton, T., Chakalova, L., Cope, N.F., Horton, A., Andrews, S., Kurukuti, S., Mitchell, J.A., Umlauf, D., Dimitrova, D.S. et al. (2009) Preferential associations between co-regulated genes reveal a transcriptional interactome in erythroid cells. Nat. Genet., 42, 53–61.

26. Babaei, S., Mahfouz, A., Hulsman, M., Lelieveldt, B.P.F., de Ridder, J. and Reinders, M. (2015) Hi-C Chromatin Interaction Networks Predict Co-expression in the Mouse Cortex. PLoS Comput. Biol., 11, e1004221.

27. Pierson, E., Consortium, G.T., Koller, D., Battle, A., Mostafavi, S., Ardlie, K.G., Getz, G., Wright, F.A., Kellis, M., Volpi, S. et al. (2015) Sharing and Specificity of Co-expression Networks across 35 Human Tissues. PLoS Comput. Biol., 11, e1004220.

28. Kulaeva, O.I., Nizovtseva, E.V., Polikanov, Y.S., Ulianov, S.V. and Studitsky, V.M. (2012) Distant activation of transcription: mechanisms of enhancer action. Mol. Cell. Biol., 32, 4892–4897.

29. Lobanenkov, V.V., Nicolas, R.H., Adler, V.V., Paterson, H., Klenova, E.M., Polotskaja, A.V. and Goodwin, G.H. (1990) A novel sequence-specific DNA binding protein which interacts with three regularly spaced direct repeats of the CCCTC-motif in the 5’-flanking sequence of the chicken c-myc gene. Oncogene, 5, 1743–1753.

30. Shlyueva, D., Stampfel, G. and Stark, A. (2014) Transcriptional enhancers: from properties to genome-wide predictions. Nat. Rev. Genet., 15, 272–286.

31. Zhu, Y., Chen, Z., Zhang, K., Wang, M., Medovoy, D., Whitaker, J.W., Ding, B., Li, N., Zheng, L. and Wang, W. (2016) Constructing 3D interaction maps from 1D epigenomes. Nat. Commun., 7, 10812.

32. Papantonis, A. and Cook, P.R. (2013) Transcription Factories: Genome Organization and Gene Regulation. Chem Rev, 113, 8683–8705.

33. Gligorijević, V. and Pržulj, N. (2015) Methods for biological data integration: perspectives and challenges. J. R. Soc. Interface, 12.

34. Gligorijević, V. and Pržulj, N. (2016), Europeanization and Globalization, pp. 137–178.

35. Lee, D.D. and Seung, H.S. (1999) Learning the parts of objects by non-negative matrix factorization. Nature, 401, 788–791.

36. Marbach, D., Lamparter, D., Quon, G., Kellis, M., Kutalik, Z. and Bergmann, S. (2016) Tissue-specific regulatory circuits reveal variable modular perturbations across complex diseases. Nat Methods, 13, 366–370.

37. Hutchins, L.N., Murphy, S.M., Singh, P. and Graber, J.H. (2008) Position-dependent motif characterization using non-negative matrix factorization. Bioinformatics, 24, 2684–2690.

